# Genomics of Ocular *Chlamydia trachomatis* after 5 years of SAFE interventions for trachoma in Amhara, Ethiopia

**DOI:** 10.1101/2020.06.07.138982

**Authors:** Harry Pickering, Ambahun Chernet, Eshetu Sata, Mulat Zerihun, Charlotte A. Williams, Judith Breuer, Andrew W. Nute, Mahiteme Haile, Taye Zeru, Zerihun Tadesse, Robin L. Bailey, E. Kelly Callahan, Martin J. Holland, Scott D. Nash

## Abstract

**Background:** To eliminate trachoma as a public health problem, the WHO recommends the SAFE strategy. As part of the SAFE strategy in the Amhara Region, Ethiopia, the Trachoma Control Program distributed over 124 million doses of antibiotic between 2007 and 2015. Despite these interventions, trachoma remained hyperendemic in many districts and a considerable level of *Chlamydia trachomatis* (*Ct*) infection was evident.

**Methods:** We utilised residual material from Abbott m2000 *Ct* diagnostic tests to sequence 99 ocular *Ct* samples from Amhara and investigated the role of *Ct* genomic variation in the continued transmission of *Ct* following 5 years of SAFE.

**Findings:** Sequences were typical of ocular *Ct*, at the whole-genome level and in tissue tropism-associated genes. There was no evidence of macrolide-resistance in this *Ct* population. Polymorphism in a region around *ompA* gene was associated with village-level TF prevalence. Additionally, greater *ompA* diversity at the district-level was associated with increased *Ct* infection prevalence.

**Interpretation:** We found no evidence for *Ct* genomic variation contributing to continued transmission of *Ct* after treatment, adding to previous evidence that azithromycin does not drive acquisition of macrolide resistance alleles in *Ct*. Increased *Ct* infection in villages and in districts with more *ompA* variants requires longitudinal investigation to understand what impact this may have on treatment success and host immunity.

**Funding:** European Commission; Neglected Tropical Disease Support Center; International Trachoma Initiative

## Introduction

Trachoma is a blinding disease caused by the intracellular bacterium *Chlamydia trachomatis* (*Ct*). To eliminate trachoma as a public health problem, the World Health Organization (WHO) recommends the SAFE (Surgery, Antibiotics, Facial cleanliness, and Environmental improvement) strategy.^1^ As part of this strategy, annual mass drug administration (MDA) with azithromycin is delivered to individuals aged ≥6 months whilst topical tetracycline eye ointment is administered to pregnant women and children aged <6 months. The number of recommended years of SAFE interventions is based on the prevalence of trachoma in a district, determined by population based surveys.^2^ For districts considered hyperendemic for trachoma, defined as a trachomatous inflammation-follicular (TF) prevalence of ≥ 30% among children aged 1 to 9 years, 5 to 7 years of SAFE are recommended followed by further population based surveys to determine the impact of the interventions.

As part of the SAFE strategy in Amhara National Regional State, Ethiopia, the Trachoma Control Program distributed over 124 million doses of antibiotic between 2007 and 2015.^3^ Both administrative and self-reported coverage have demonstrated treatment coverage in the region to be close to or above the WHO recommended minimum threshold of 80%.^3–5^ During this time, the program also provided village-based and school-based health education and assisted in the construction of latrines as part of the F and E components of SAFE.^3^ Despite an average of 5 years of these interventions, trachoma remained hyperendemic in many districts, and a considerable level of *Ct* infection was evident throughout the region.^3,6^

Historically, *Ct* molecular epidemiology predominantly focused on *ompA*,^7^ the gene encoding the major outer membrane protein. More recently a number of multilocus sequence typing schemes with and without *ompA* have been used.^8–10^ Since 2010, there has been a rapid expansion of *Ct* whole-genome sequencing (WGS), due to the ability to sequence directly from clinical samples.^11,12^ Despite more than seven-hundred *Ct* genomes being publicly available,^13–16^ few studies have evaluated the role of genome-level variation in *Ct* transmission and clinical outcomes of *Ct* infection. Recent publications have begun to address these questions in *Ct* populations from trachoma-endemic settings^16,17^ and have shown higher diversity than expected, compared with sequencing of cultured isolates. WGS additionally allows monitoring of emergence of antimicrobial resistance alleles in *Ct*,^17–19^ which is of critical importance as MDA with azithromycin is key for trachoma control, and is also under consideration as an intervention for childhood mortality,^20,21^ neonatal sepsis,^22^ and bacterial skin diseases.^23^

The Trachoma Control Program in Amhara has conducted multiple studies to better understand the epidemiology of trachoma in communities that have received approximately 5 years of annual MDA with azithromycin, yet still have significant levels of *Ct* infection and trachomatous disease. The resolution of WGS allows for a greater understanding of *Ct* transmission patterns, presence of putative virulence determinants, and identification of antimicrobial resistance alleles. Therefore, this study sequenced 99 ocular *Ct* samples from Amhara to investigate the role of genomic variation in the continued transmission of *Ct*. We further explored the relationship between *Ct* genomic variation, ocular *Ct* infection prevalence, and trachoma clinical sign prevalence at the village and district level.

## Methods

### Ethics statement

Survey methods were approved by the Emory University Institutional Review Board (protocol 079-2006) as well as the Amhara Regional Health Bureau. Due to the high illiteracy rate among the population, approval was obtained for oral consent or assent. Further permission for sample transfer and genomic sequencing of *Ct* samples for this project was provided by the Amhara Regional Health Bureau.

### Study design and population

Between 2007 and 2010 the SAFE strategy was scaled to reach all districts in Amhara and interventions were subsequently administered for approximately 5 years. Sampling methodology for these district-level surveys has been published previously.^3,24^ Briefly, a multi-stage cluster randomized methodology was used, whereby clusters (villages) were selected using a population proportional to estimated size method, and within a cluster, a modified segmentation approach was used to randomly select segments of 30 to 40 households.^3^

After enumerating all residents, present and consented residents were examined for trachoma based on the WHO simplified methodology.^25^ Trained and certified trachoma graders used x2·5 magnification and adequate light.^3,26^ Every-other cluster was chosen for swab collection prior to surveying a district, and during the house-to-house survey, the first 25 children aged 1 to 5 years with parental consent were swabbed for the presence of infection. If more than one child aged 1 to 5 years lived in a household, one child was randomly chosen by survey software.

### Sample collection and processing

Gloved graders swabbed the upper tarsal conjunctiva three times with a polyester-tipped swab, rotating 120 degrees along the swab’s axis each time to collect a sufficient epithelial specimen.^6^ Samples were transferred to the Amhara Public Health Institute (APHI) where they were stored at −20° C. Conjunctival swabs from each district were randomized and five samples were combined into each pool. Pools were processed with the RealTime (Abbott Molecular Inc., Des Plaines, IL, USA) polymerase chain reaction assay on the Abbott m2000 system at the APHI laboratory.^6^ All individual samples from the identified positive pools from four zones (administrative groupings of districts), North Gondar, South Gondar, East Gojam, and Waghemra, were processed again to provide individual level infection data.^27^ Samples from these zones were prioritized owing to the persistent high trachoma prevalence. For positive individual samples, the m2000 generated delta cycle threshold result was converted to *Ct* elementary body (EB) equivalent concentration based on a previously described calibration curve of known EB concentrations on the RealTime Assay.^27,28^

Once *Ct* load was known for the positive individual samples, a total of 240 with the highest load were chosen for this project. Samples with sufficient *Ct* load, likely to obtain high quality full genome sequence data based on our previous studies, were re-extracted as described below.^16,17,29^

### Ct detection and sequencing preparation

DNA was extracted from 800 μl residual material per sample from Abbott m2000 diagnostic tests using QIAamp mini DNA kit (Qiagen). Samples were quantified using plasmid and genome targets by quantitative PCR as previously described.^30^ Samples with ≥ 10 genome copies per μl of DNA were considered for WGS.

### Sequencing, processing, and analysis of Ct

Sequencing was performed as previously described^16^ except we utilised the SureSelectXT Low Input kit. Processing and analysis of sequenced reads was performed as previously described.^17^ Briefly, raw reads were trimmed and filtered using Trimmomatic.^31^ Filtered reads were aligned to a reference genome (A/Har13) with Bowtie2,^32^ variant calls were identified with SAMtools/BCFtools.^33^ Multiple genome and plasmid alignments were generated using progressiveMauve,^34^ multiple gene alignments were generated using muscle.^35^ Phylogenies were computed using RaxML,^36^ predicted regions of recombination were masked using Gubbins^37^ before visualisation in R. Domain structure of *tarP* and truncation of *trpA* were characterised as previously described.^16^ ABRicate and the ResFinder database (https://github.com/tseemann/abricate) were used to test for the presence of antimicrobial resistance genes in the reference-assembled genomes and de novo assembled reads for each sample.

### Inter-population genome-wide association analysis

Genome-wide association analysis was performed to identify polymorphisms specific to this population of ocular *Ct* sequences through comparison of 99 Amharan *Ct* genomes to 213 previously sequenced samples from trachoma-endemic communities. Heterozygous calls and positions with greater than 25% missing data were removed. Polymorphisms were considered conserved in Amhara if the major allele frequency was greater than 0.8 and rare in the global ocular population if the same allele was at a frequency of less than 0.2. The final analysis included 116 single nucleotide polymorphisms (SNPs). A logistic regression was performed with each Amhara-specific site as the independent variable and origin of the sequence as the dependent variable (reference level; global and comparator level; Amharan). P-values were considered for significance after Bonferroni correction.

### Intra-population genome-wide association analyses

Genome-wide association analyses were performed to identify *Ct* polymorphisms associated with village-level clinical data. Heterozygous base calls were and positions with a minor allele frequency of less than 25% or greater than 25% missing data were removed. The final analysis included 681 SNPs across the 99 sequences. A linear regression was performed with each SNP as the independent variable and log_10_ transformed village-level *Ct* infection, TF, or TI prevalence as the dependent variable. District was included as a random-effect and all analyses were adjusted for age and gender. P-values were considered significant after Bonferroni correction. Additionally, a sliding-window approach was used to identify polymorphic regions of the genome. Windows of 10 kilobases were evaluated for polymorphic sites, with a step size of 1 kilobase. There were a median of four polymorphic sites per window (IQR; 2-7). The final analysis included 907 polymorphic regions across the 99 sequences. A linear regression was performed with each polymorphic region collapsed into a pseudo-haplotype per sequence as the independent variable, including district as a random-effect and adjusted for age and gender. This model was compared to a model including only the covariates and random-effects by F-test. P-values were considered significant after Bonferroni correction.

### Inference of ompA sequences

Complete sequences of *ompA* were obtained from whole-genome sequence data using the reference-based assembly method described above with one change. Each sample was assembled against four reference sequences (A/Har-13, B/Jali-20, C/TW-3, and D/UW-3) and the assembly with the highest coverage was used for downstream analyses. Serovar of *ompA* was assigned using maximum *blastn* homology against all published *Ct* sequences. Genotypes of *ompA* were manually determined using SeaView.^38^ Diversity of *ompA* genotypes was calculated as Simpson’s D using *vegan* in R.

## Results

Ocular swabs previously confirmed as positive for *Ct* DNA using the Abbott m2000 system were selected for this study (n = 240). One-hundred and thirty-five samples had sufficiently high concentration of *Ct* DNA after reextraction to be considered for WGS (≥ 10 genome copies per μl). Of these, 99 were randomly selected for sequencing to match the complete dataset on age, gender, and zone of collection. The sequenced and complete samples were comparable (Table 1), except as expected a higher median load of infection in sequenced samples.

**Table 1.**
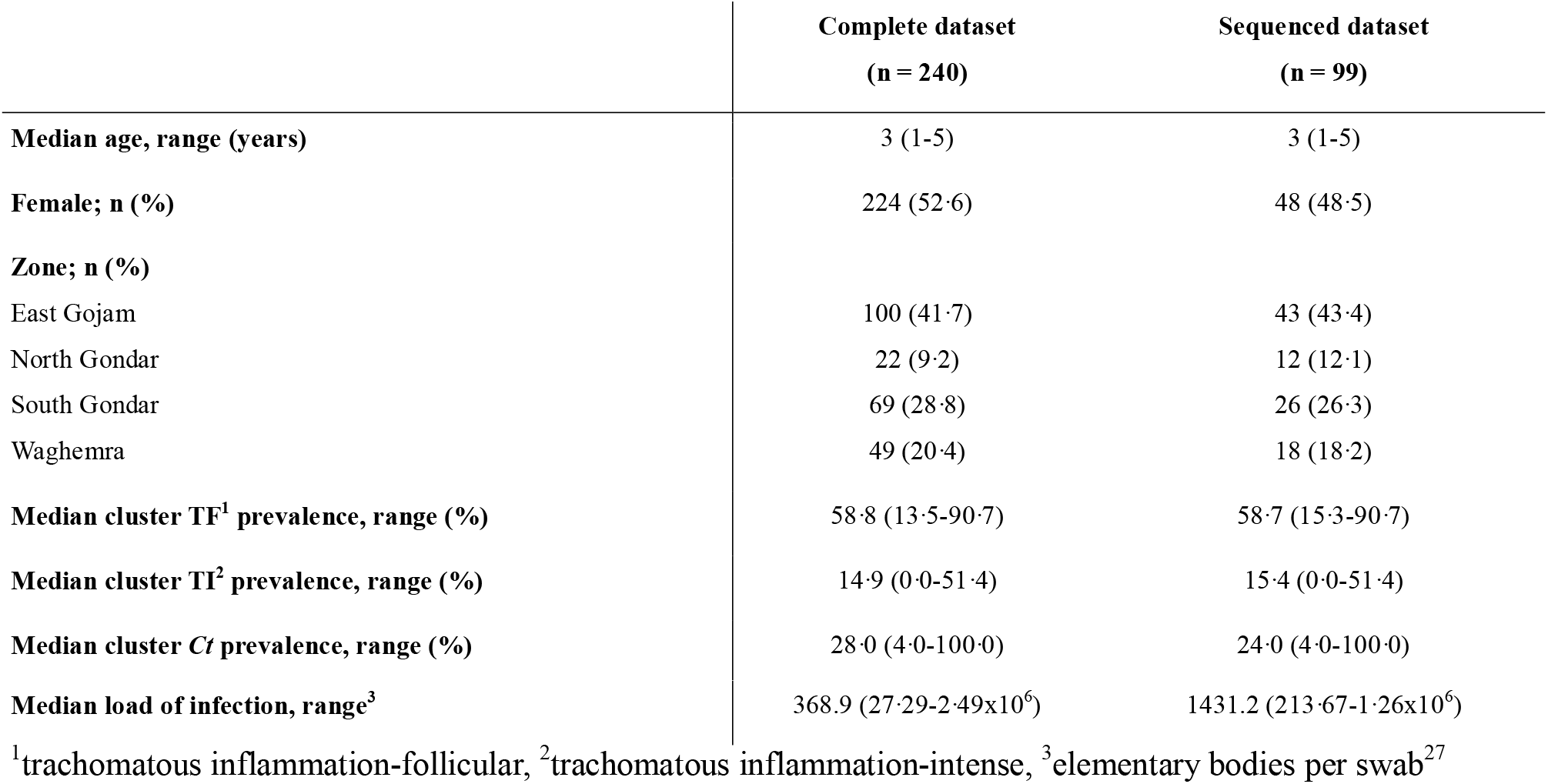
Demographic and trachoma characteristics of complete and sequenced samples, Amhara, Ethiopia, 2011-2015.

The Amharan *Ct* genomes formed two closely related subclades within the classical T2 ocular clade (Figure 1). The two subclades were predominantly separated by *ompA* genotype, with 52 serovar A (SvA) and 47 serovar B (SvB) genomes. Focusing on genomes from ocular infections (Figure 2), the SvA Amharan genomes branch together independent from any previously sequenced *Ct*. The SvB Amharan genomes were split across two branches. One branch was most closely related to A/Har-13 originally isolated from Saudi Arabia. The second, smaller branch was most closely related to Ba/Apache-2 *Ct* from the United States of America (USA) as well as recently sequenced ocular *Ct* from Solomon Islands.

**Figure 1.**
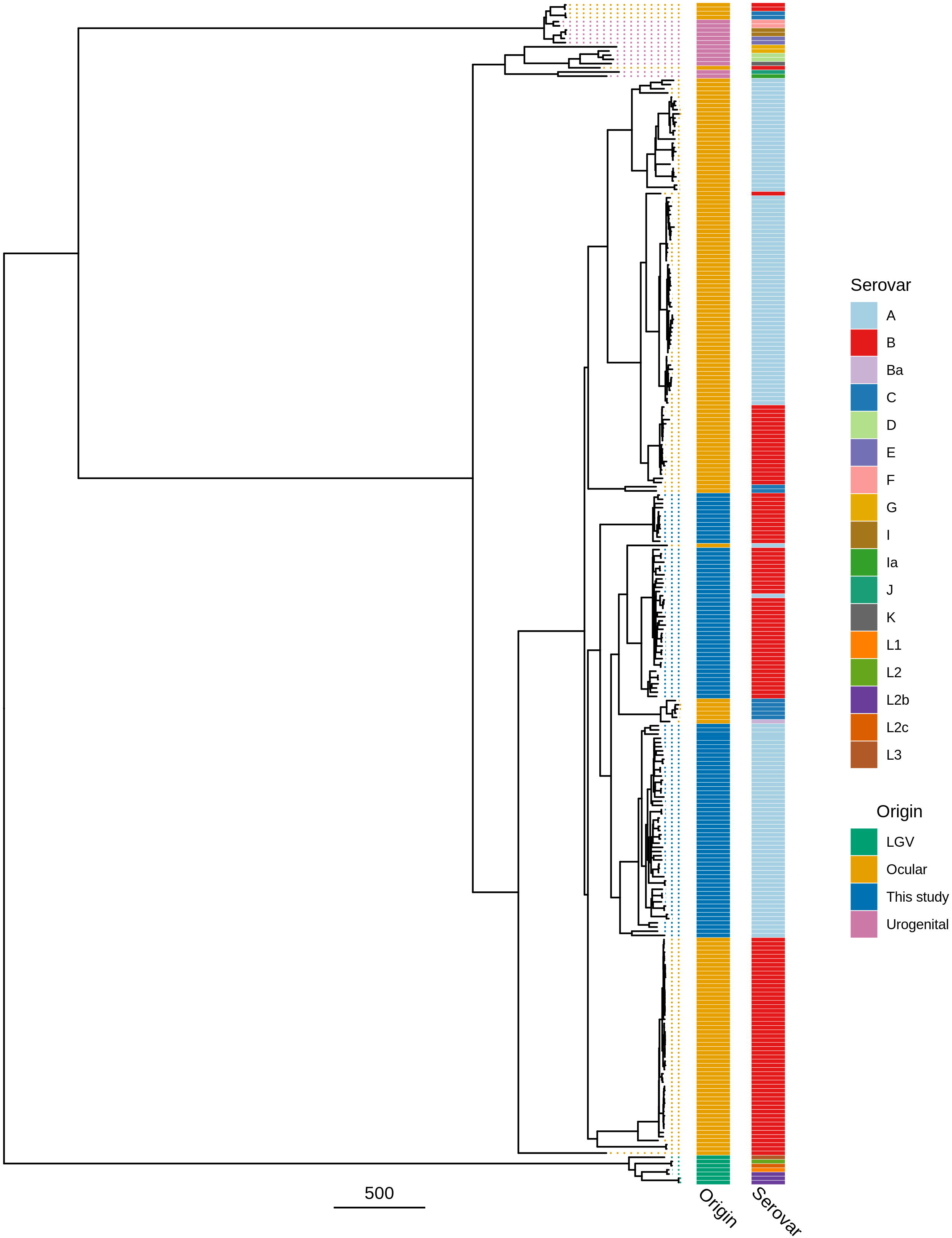
Maximum likelihood reconstruction of whole genome phylogeny of ocular Chlamydia trachomatis (Ct) sequences from Amhara, Ethiopia. Whole genome phylogeny of 99 *Ct* sequences from Amhara and 183 *Ct* clinical and reference strains. Amharan *Ct* sequences were mapped to *Ct* A/HAR-13 using Bowtie2. SNPs were called using SAMtools/BCFtools. Phylogenies were computed with RAxML from a variable sites alignment using a GTR + gamma model and are midpoint rooted. The scale-bar indicates pairwise distance. *Ct* sequences are coloured by origin of the sample (“Origin”) and *ompA* serovar (“Serovar”).

**Figure 2.**
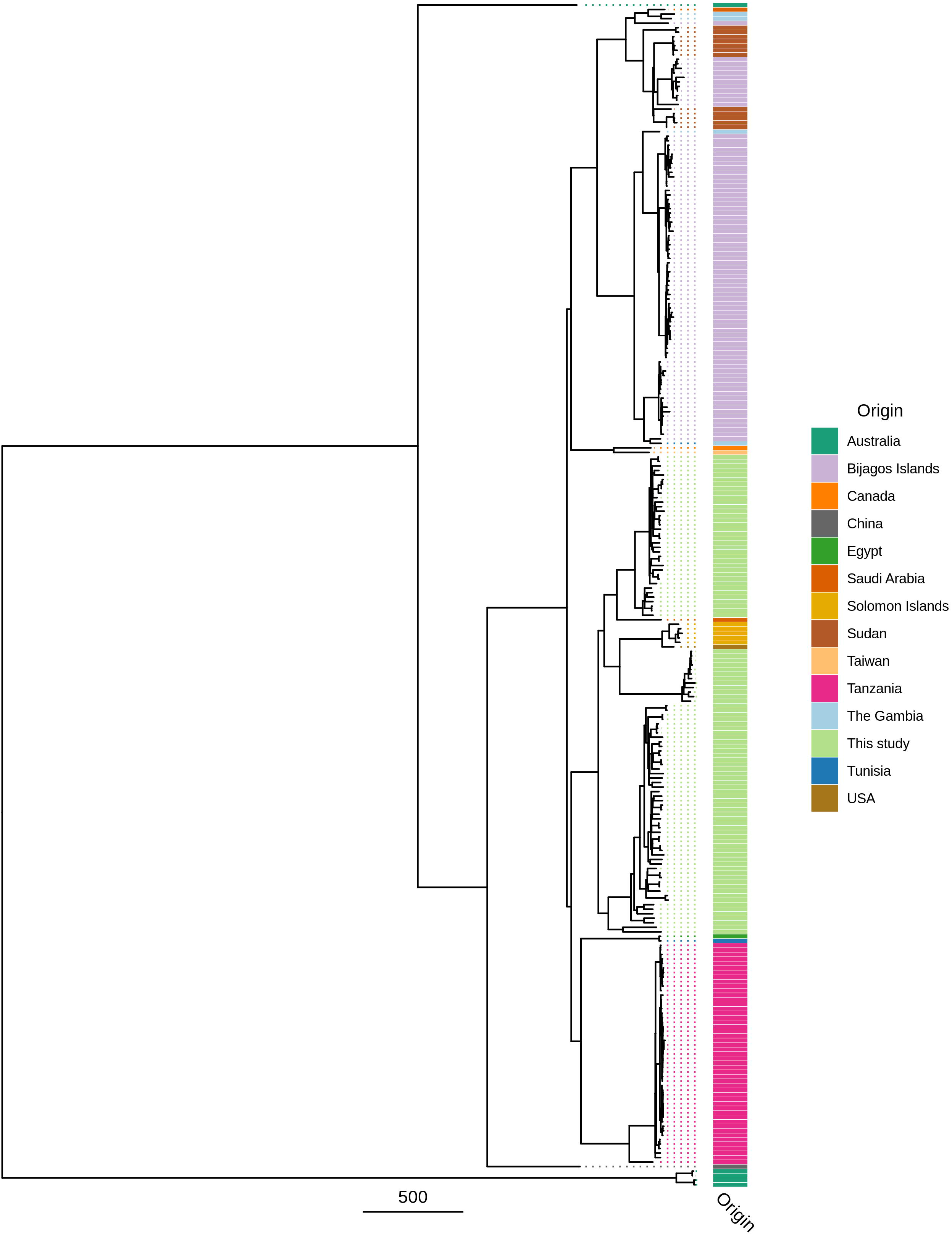
Maximum likelihood reconstruction of *ompA* phylogeny of ocular Chlamydia trachomatis (Ct) sequences from Amhara, Ethiopia. Phylogeny of *ompA* from 99 *Ct* sequences from Amhara and 183 *Ct* clinical and reference strains. Amharan *Ct* sequences were mapped to *Ct* A/HAR-13 using Bowtie2. SNPs were called using SAMtools/BCFtools. Phylogenies were computed with RAxML from a variable sites alignment using a GTR + gamma model and are midpoint rooted. The scale-bar indicates pairwise distance. *Ct* sequences are coloured by country of origin of the sample (“Origin”).

Several *Ct* genes and genomic regions are hypothesised to be indicative of tissue preference/tropism, with polymorphisms distinct to ocular, urogenital and LGV sequences. All Amharan *Ct* genomes had *tarP* domain structure typical of ocular sequences.^39^ Similarly, all Amharan genomes had inactivating mutations in *trpA*, leading to a nonfunctional tryptophan synthase.^40^ Polymorphic membrane proteins (*pmp*) *B, C, F, G, H* and *I* clustered phylogenetically with ocular isolates (SI Figure 1).^41^ There was minimal polymorphism in the *Ct* plasmid within the Amharan genomes and they were closely related to previously sequenced ocular isolates (SI Figure 2). There was no evidence for the presence of macrolide resistance alleles in the assembled genome sequences or de novo assembled sequence reads.

Amharan *Ct* genomes were compared to 213 previously sequenced samples from trachoma-endemic communities to identify genomic markers specific to Amhara.^13–17,29^ Of 36,805 polymorphic sites (Figure 3a), 116 were conserved in Amhara (frequency ≥ 0.8) and rare in the global ocular population (frequency ≤ 0.2). These were dispersed throughout the genome (Figure 3b). Fourteen genes harboured two such sites and five genes contained three sites, all of which have previously been identified as polymorphic in distinct populations of *Ct* genomes (Figure 3c).

**Figure 3.**
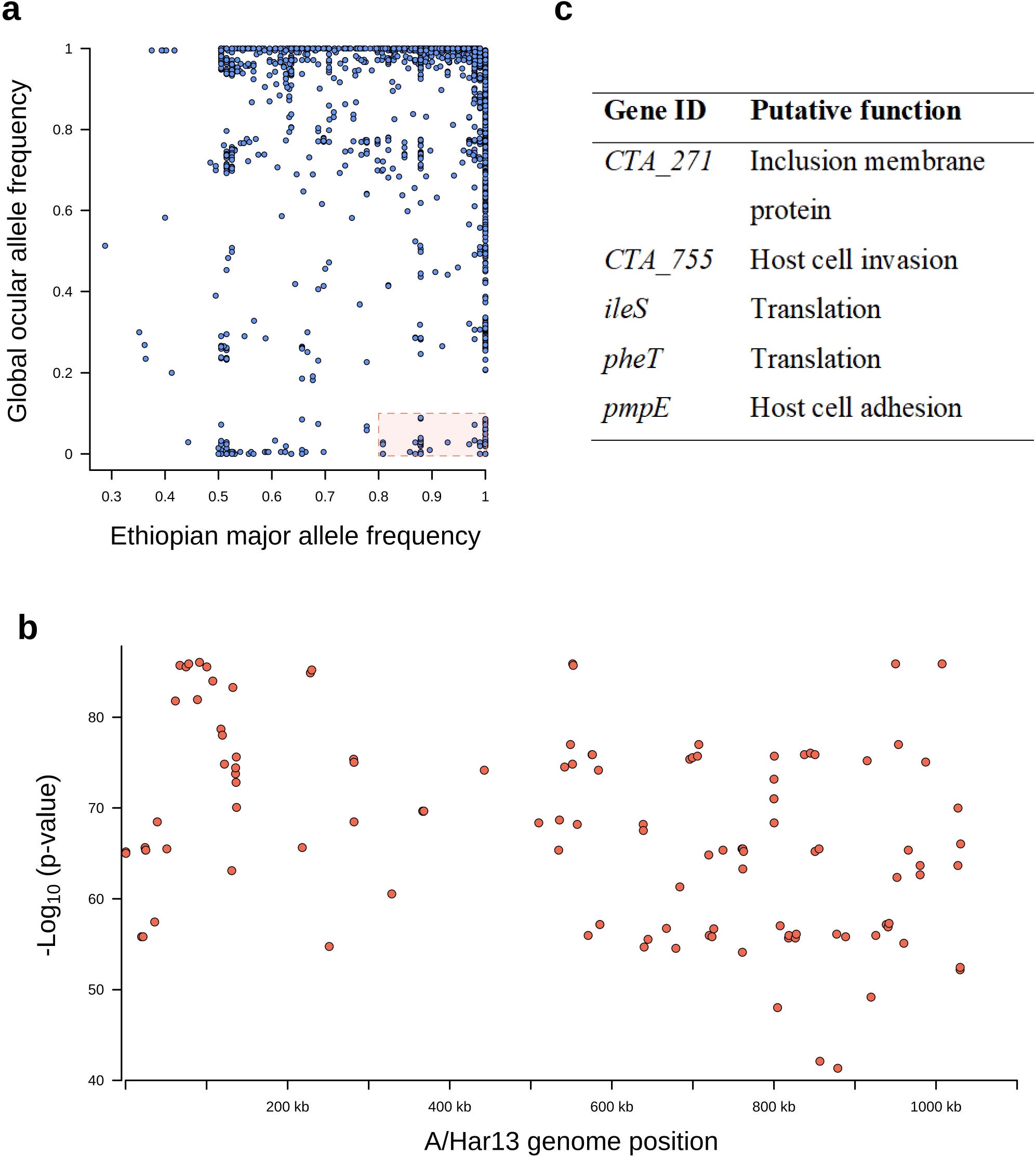
Single nucleotide polymorphisms on the Chlamydia trachomatis (Ct) genome specific to Amhara, Ethiopia. a) Single nucleotide polymorphisms (SNPs) conserved in Amhara, Ethiopia (allele frequency □≥□0.8) and rare in other *Ct* sequences (allele frequency ≤□0.2) were identified by comparing these *Ct* sequences (n□=□99) to ocular genomes from other populations (n□=□213). b) Logistic regression found SNPs specific to this Amharan population to be dispersed throughout the genome (n = 116). c) Five genes harboured three Amhara-specific SNPs, putative function was determined by reference to published literature.

A genome-wide association study was performed to identify polymorphism within the Amharan *Ct* genomes related to village-level prevalence of *Ct* infection. The final analysis included 681 single nucleotide polymorphisms (SNPs) in 99 genomes. No SNPs were associated with village-level prevalence of infection (SI Figure 3a). A secondary sliding-window approach was utilised to identify polymorphic regions of the genome associated with infection prevalence. The final analysis included 907 polymorphic regions in 99 genomes. No polymorphic regions were associated with village-level prevalence of infection (SI Figure 3b).

No SNPs were associated with village-level prevalence of TF (Figure 4a). However, eight polymorphic regions from positions 774,000 to 791,000 were associated with village-level prevalence of TF (Figure 4b). SNPs in these regions were focused in CTA0743/*pbpB* (harbouring 29 SNPs), *CTA0747/sufD* (10 SNPs) and CTA0742/*ompA* (7 SNPs). All SNPs in *sufD* were synonymous, while 8/29 and 3/7 SNPs in *pbpB* and *ompA* were non-synonymous.

**Figure 4.**
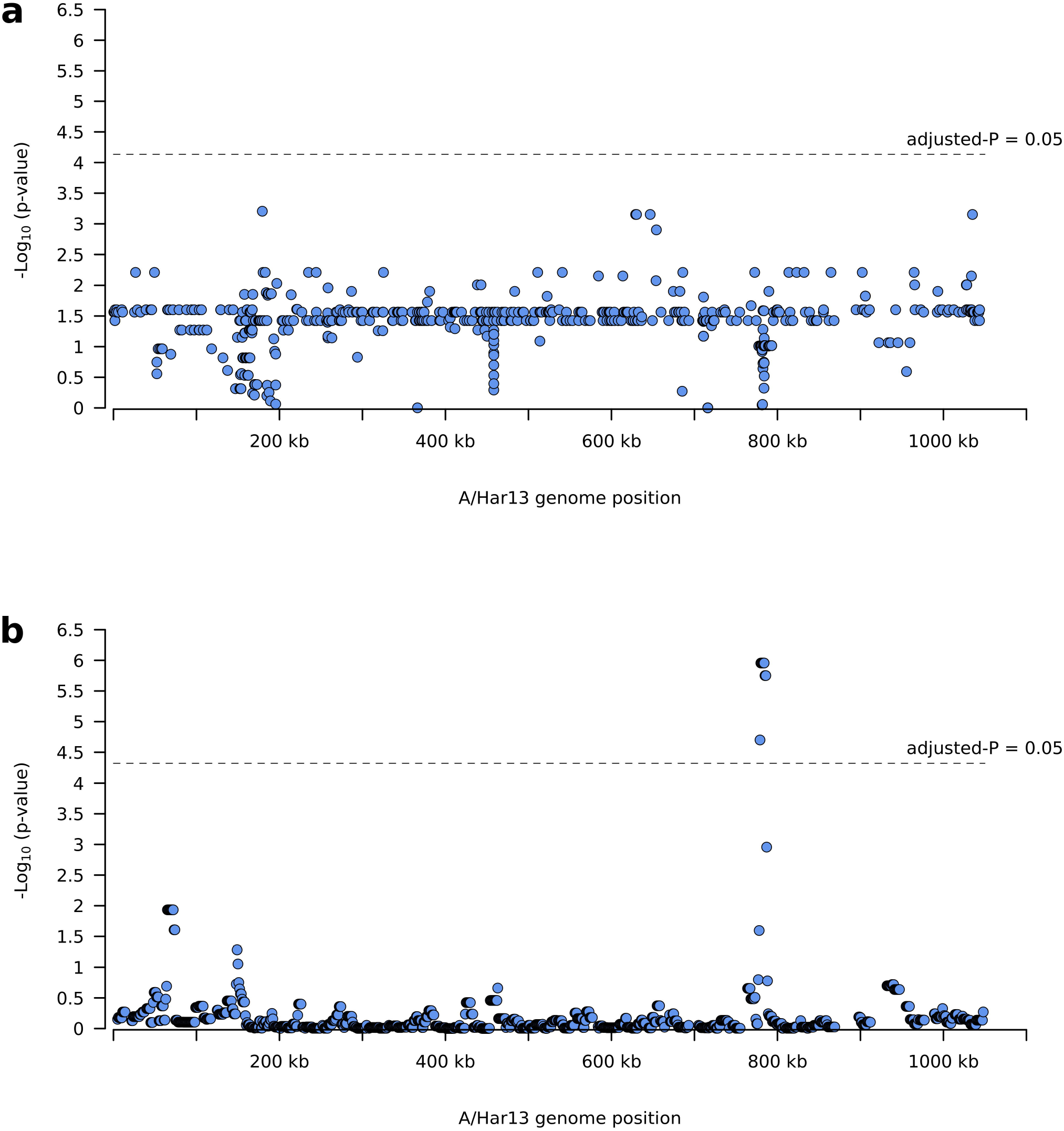
Polymorphisms on the *Chlamydia trachomatis* (*Ct*) genome associated with village-level TF prevalence. a) No single nucleotide polymorphisms were significantly associated with village-level TF prevalence. b) Eight polymorphic regions from positions 774,000 to 791,000 were associated with village-level prevalence of TF.

No SNPs or polymorphic regions were associated with village-level prevalence of TI (SI Figure 4).

As *ompA* variation was important in *Ct* phylogeny and heterogeneity in TF profiles in this population, we further investigated the geographical distribution of *ompA* serovars and their relationship to levels of *Ct* infection and TF. Serovars A (SvA) and B (SvB) of *ompA* were distributed across all studied zones (Figure 5a). Village-level *Ct* infection, TF and TI prevalence were not associated with *ompA* serovar (p = 0·860, 0·382 and 0·177 respectively). We identified nine *ompA* types in this population (Table 2). Six were serovar A (SvA), defined by nine non-synonymous polymorphic sites. Three were serovar B (SvB), defined by two non-synonymous polymorphic sites. Four of nine types were present in all zones of this study (A1, A3, A5 and B3), four were exclusive to East Gojam (A2, A4, A6 and B1) and one was found in both East Gojam and North Gondar (B2) (SI Figure 5). Types A1 (n = 5) and B1 (n = 6) had a nucleotide predicted amino acid change in the surface-exposed, variable domain 1 (VD1), A2 (n = 2) in VD2 and A4 (n = 1) in VD4.

**Figure 5.**
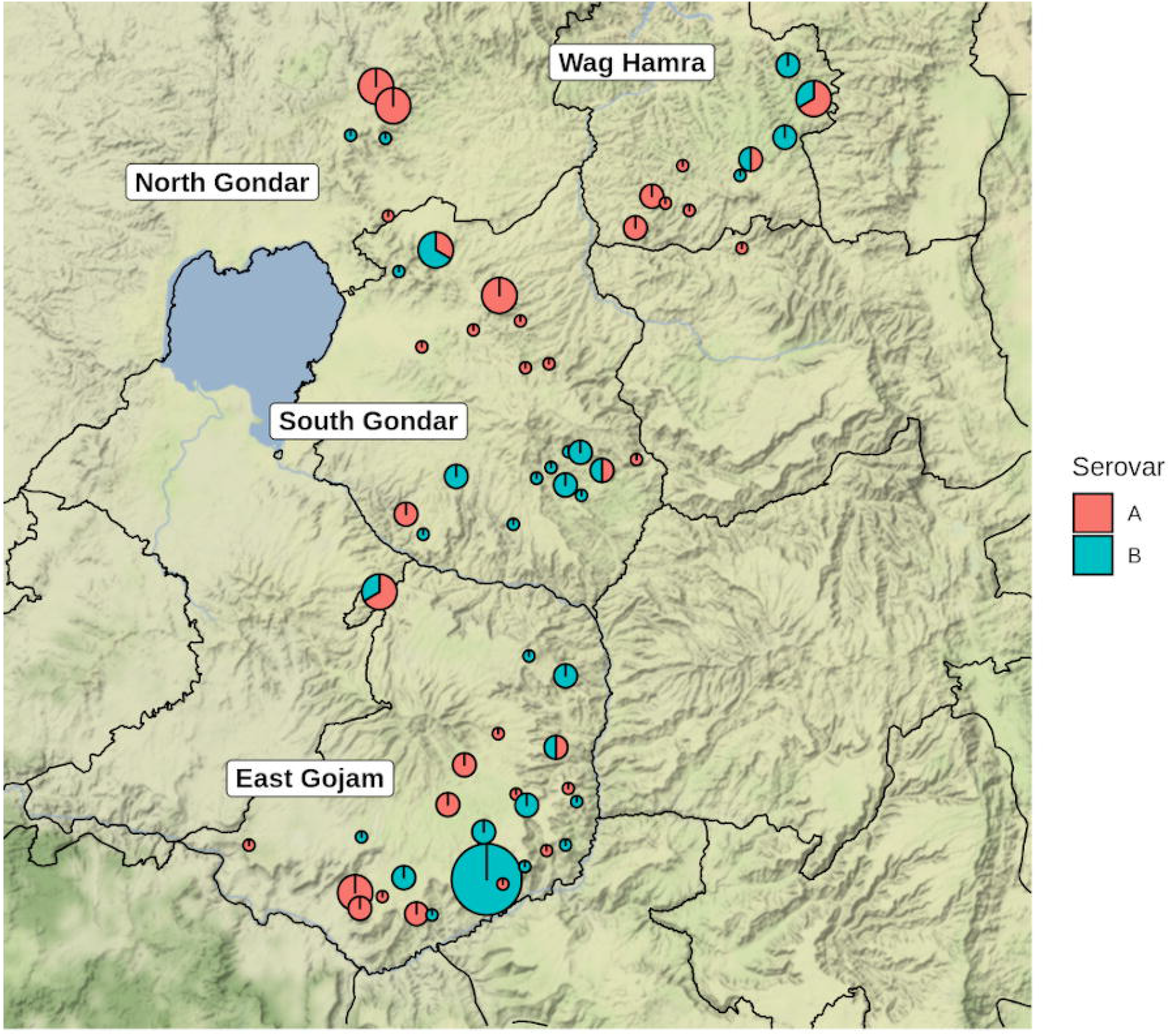
Geographical distribution and similarity of *ompA* serovars. a) Four zones in Amhara, Ethiopia were represented in this study. Pie charts represent village-level *Ct* prevalence (pie diameter) and presence of *ompA* serovars A (red) and B (blue). Maps were generated using R package ggmap, shape files were obtained from Google Maps.

**Table 2.** Description of nucleotide polymorphisms and amino acid changes in ompA of Amharan Ct sequences.

| *ompA*<br>Type<br>(number) | Nucleotide Position and Reference Nucleotide (Amino Acid Position and Reference Amino Acid) |  |  |  |  |  |  |  |  |  |  |
| --- | --- | --- | --- | --- | --- | --- | --- | --- | --- | --- | --- |
|  | Serovar A |  |  |  |  |  |  |  |  | Serovar B |  |
|  | 272G<br>(91S) | 305C<br>(102A) | 433A<br>(145T) | 736A<br>(246I) | 940A<br>(314K) | 943C<br>(315P) | 946G<br>(316V) | 955A<br>(319T) | 956C<br>(319T) | 286A<br>(96T) | 1132G<br>(378A) |
| A1 (5) | A<br>(Asp) |  |  | G (Ile) |  |  |  |  |  |  |  |
| A2 (2) |  |  | G (Ala) | G (Ile) |  |  |  |  |  |  |  |
| A3 (21) |  |  |  | G (Ile) |  |  |  |  |  |  |  |
| A4 (1) |  |  |  |  | G (Glu) | G (Ala) | A<br>(Ile) | G (Val) | T (Val) |  |  |
| A5<sup>ref</sup> (22) |  |  |  |  |  |  |  |  |  |  |  |
| A6 (1) |  | T<br>(Val) |  |  |  |  |  |  |  |  |  |
| B1 (6) |  |  |  |  |  |  |  |  |  | G (Ala) | A<br>(Thr) |
| B2 (6) |  |  |  |  |  |  |  |  |  |  | A<br>(Thr) |
| B3<sup>ref</sup> (35) |  |  |  |  |  |  |  |  |  |  |  |

<sup>ref</sup>reference type per serovar to determine classify variants

Most villages (55/61) had only one *ompA* type in this study, therefore we evaluated *ompA* diversity at the district-level, using Simpson’s D. We used previously published district-level *Ct* infection, TF and TI prevalence estimates^6,26^ (Table 3). District-level *Ct* infection and TI prevalence were significantly higher with increasing *ompA* diversity, a similar trend was found for TF prevalence. In a multivariate model, only *Ct* infection prevalence was associated with increasing *ompA* diversity.

**Table 3.** Linear regression analysis of predictors of district-level *ompA* diversity.

| Variable<sup>1</sup> | Univariate |  |  | Multivariate |  |  |
| --- | --- | --- | --- | --- | --- | --- |
| | $\beta^2$ | SE<sup>3</sup> | p-value | $\beta$ | SE | p-value |
| ***Ct* infection prevalence** | 1.134 | 0.238 | 0.69x10<sup>-4</sup> | 0.402 | 0.114 | 0.002 |
| **TF prevalence** | 0.005 | 0.004 | 0.159 | -0.004 | 0.004 | 0.275 |
| **TI prevalence** | 0.033 | 0.012 | 0.008 | 0.016 | 0.013 | 0.260 |

<sup>1</sup>district-levels prevalence estimates, <sup>2</sup> $\beta$ = regression coefficient, <sup>3</sup>SE = standard error

## Discussion

This study sequenced *Ct* from ocular samples collected from districts in Amhara, Ethiopia which had received approximately 5 years of the SAFE strategy, including annual azithromycin MDA, as part of trachoma control efforts. We found sequences were typical of ocular *Ct*, at both the whole-genome level and in tropism-associated genes, yet phylogenetically distinct from most previously sequenced *Ct* genomes. There was no evidence of macrolide-resistance alleles in this ocular *Ct* population. Greater *ompA* diversity at the district-level was associated with increased *Ct* infection prevalence. A continued commitment to the implementation of the full SAFE strategy with consideration of enhanced MDA accompanied by further longitudinal investigation is warranted in Amhara.

Almost 900 million doses of azithromycin have been distributed by trachoma control programmes since 1999^42^ and in Amhara alone 15 million doses are administered every year.^3^ Mass distribution of azithromycin is likely to become more common as more evidence emerges of off-target effects such as reducing infectious diseases,^23,43–45^ diarrheal diseases^46^ and childhood mortality.^22,47–50^ There is concern about the impact of these programmes on development of antimicrobial resistance in *Ct* and other bacteria.^19,51^ This is particularly true in situations where community-wide treatment with azithromycin has been unable to eliminate trachoma as a public health problem within expected timelines.^46,52^ It has been shown that treating whole communities with azithromycin can increase nasopharyngeal carriage of macrolide-resistant *Staphylococcus*^53^ and *Streptococcus*^54,55^ species and alters the faecal microbiome,^56–59^ with studies reporting an increase in macrolide-resistant *Escherichia coli*.^60,61^ This study, in agreement with previous work,^17–19^ found no evidence of macrolide-resistance alleles in this *Ct* population. This result is encouraging; however, it does not rule out macrolide-resistance as a potential problem in these communities. Carriage of macrolide-resistant pathogens in the gut and nasopharynx may be impacted by continued antibiotic treatment. Additionally, presence of additional species of *Chlamydia*^62,63^ and non-chlamydial bacteria^64–69^ in the ocular niche have been associated with clinical signs of trachoma, therefore resistance in other bacteria may be important.

No *Ct* genomes in this study had acquired azithromycin resistance alleles, however, there may be other genomic factors which support *Ct* transmission after treatment. To answer this question, we compared the Amharan *Ct* genomes with previously sequenced *Ct* to find polymorphism specific to this population that could explain continued transmission. The few SNPs identified as conserved in Amhara and rare in the global ocular *Ct* population were dispersed across the genome in known polymorphic genes, rather than being overrepresented in genes related to *Ct* survival. The typical nature of this *Ct* population was further supported by phylogenetic clustering with other ocular *Ct* sequences, presence of a non-functional tryptophan synthase operon and tropism-associated polymorphism in *tarP* and the polymorphic membrane proteins. Similarly to recent studies from distinct trachoma-endemic communities,^13–17,29^ the *Ct* sequences in this population formed two closely related subclades within the ocular clade, primarily separated by *ompA* serovar. Evidence of phylogenetic clustering by country of collection and the similarity to *Ct* sequences collected over 50 years prior to this study suggests diversification in ocular *Ct* is slow and geography-related, rather than being driven by treatment-derived selection pressure. A surprising finding in this study was that a subgroup of SvB *Ct* from Amhara were most closely related to a historical genome from USA (Ba/Apache-2) and recently collected genomes from Solomon Islands.^29^ It is possible the origin of these genomes is unique within this population; however, it is more likely that this is further evidence of the slow diversification of *Ct*. In support of this, *ompA* SvB sequences were significantly less diverse than SvA in this study. Furthermore, all major branches of ocular *Ct* phylogeny studied here included samples collected decades apart from geographically disparate sites.

We identified several polymorphic regions associated with village-level TF prevalence, a similar trend was found with village-level *Ct* and TI prevalence. The polymorphisms were mostly frequently found in *ompA*, *pbpB* and *sufD*, all of which are known to be polymorphic in ocular Ct.^8,70–72^ *OmpA* encodes the major outer membrane protein which constitutes approximately 2/3 of the surface of *Ct*, is the primary target of host immune responses and is believed to function as a porin.^73^ The functions of *pbpB* and *sufD* in *Ct* are currently unknown,^74^ however bacterial homologues of these genes function in peptidoglycan synthesis and response to oxidative stress^75^ respectively. It is plausible that genes hypothesised to be involved in immune evasion and response to stress could have an impact on *Ct* survival and response to treatment.

We found approximately equal representation of SvA (52.5%) and SvB (47.5%) in this sample of the Amhara population. Both serovars were present in all four districts and were not individually associated with village-level *Ct* infection, TF, or TI prevalence. However, *Ct* infection prevalence, and to a lesser extent TF and TI prevalence, were increased in districts with greater *ompA* diversity. Our data agrees with that from a Nepalese study that found increased *ompA* diversity in villages to be associated with higher trachoma prevalence.^76^ In contrast, a more recent study from Ethiopia found no association between *ompA* diversity and *Ct* infection levels^77^. It is known that individuals develop serovar-specific immunity to *Ct*,^78–80^ therefore it is plausible that in villages with more than one serovar in circulation, individuals are more likely to be exposed to a serovar they do not have protective immunity against. However, presence of one or more *ompA* variants should not impact treatment success.

A potential limitation of this study was bias towards samples with the highest *Ct* load. It is possible that identified relationships between *ompA* variation and *Ct* infection prevalence might have been different if lower load infections were included, particularly at the village-level, as the majority (34/61) were represented by one sequence. Additionally, we have not sequenced residual material from Abbott m2000 specimens previously, therefore, it is possible that long-term storage in this format and multiple freeze-thaw cycles may have impacted DNA quality and/or quality. However, obtaining high quality genomes from all sequenced samples, with as low as 500 *Ct* genomes as starting material, suggests quality was not an issue.

Despite approximately five years of azithromycin MDA, we found no evidence for *Ct* genomic variation contributing to the continued transmission of *Ct* after multiple rounds of treatment, adding to previous evidence that azithromycin MDA does not drive acquisition of macrolide resistance alleles in *Ct*. This study demonstrates feasibility of WGS of low-load, residual material and highlights the added value of collecting ocular swabs as part of routine trachoma surveys. Collection and long-term storage of these samples has helped alleviate concerns of azithromycin resistance in Amharan *Ct*, while offering important insights into the relationship between *ompA* variation and *Ct* infection levels. Future longitudinal investigation will be needed to understand what impact, if any, ompA diversity may have on treatment success in this setting.

## Contributors

HP, RLB, EKC, MJH and SDN contributed to study design. HP, CAW, AC, ES, MZ, ZT, EKC, and SDN contributed to data collection. HP, AWN, EKC, MJH and SDN contributed to data analysis. All authors interpreted the findings, contributed to writing the manuscript, and approved the final version for publication.

## Acknowledgments

The authors would like to acknowledge the study participants and field team in Amhara, Ethiopia. The authors also acknowledge the infrastructure support provided by the UCL/UCLH Biomedical Research Centre funded Pathogen Genomics Unit. We would also like to thank Abbott for its donation of the m2000 RealTi*m*e molecular diagnostics system and consumables.

## Availability of data and materials

All sequence data are available from the European Bioinformatics Institute (EBI) short read archive (PRJEB38668).

## References

1. Mariotti, S. P., Pararajasegaram, R. & Resnikoff, S. Trachoma: looking forward to Global Elimination of Trachoma by 2020 (GET 2020). The American journal of tropical medicine and hygiene (2003) doi:10.4269/ajtmh.2003.69.5_suppl_1.0690033.

2. Emerson, P. M., Burton, M., Solomon, A. W., Bailey, R. & Mabey, D. The SAFE strategy for trachoma control: Using operational research for policy, planning and implementation. Bulletin of the World Health Organization (2006) doi:10.2471/BLT.05.28696.

3. Stewart, A. E. P. et al. Progress to eliminate trachoma as a public health problem in Amhara National Regional State, Ethiopia: Results of 152 population-based surveys. Am. J. Trop. Med. Hyg. (2019) doi: 10.4269/ajtmh.19-0450.

4. Astale, T. et al. Population-based coverage survey results following the mass drug administration of azithromycin for the treatment of trachoma in Amhara, Ethiopia. PLoS Negl. Trop. Dis. (2018) doi:10.1371/journal.pntd.0006270.

5. Ebert, C. D. et al. Population coverage and factors associated with participation following a mass drug administration of azithromycin for trachoma elimination in Amhara, Ethiopia. Trop. Med. Int. Heal. (2019) doi:10.1111/tmi.13208.

6. Nash, S. D. et al. Ocular Chlamydia trachomatis infection under the SAFE strategy in Amhara, Ethiopia, 2011-2015. Clin. Infect. Dis. (2018) doi:10.1093/cid/ciy377.

7. Geisler, W. M., Black, C. M., Bandea, C. I. & Morrison, S. G. Chlamydia trachomatis OmpA genotyping as a tool for studying the natural history of genital chlamydial infection. Sex. Transm. Infect. (2008) doi:10.1136/sti.2008.030825.

8. Klint, M. et al. High-resolution genotyping of Chlamydia trachomatis strains by multilocus sequence analysis. J. Clin. Microbiol. (2007) doi:10.1128/JCM.02301-06.

9. Patiño, L. H. et al. Unveiling the multilocus sequence typing (MLST) schemes and core genome phylogenies for genotyping chlamydia trachomatis. Front. Microbiol. (2018) doi:10.3389/fmicb.2018.01854.

10. Pedersen, L. N., Pødenphant, L. & Møller, J. K. Highly discriminative genotyping of Chlamydia trachomatis using omp1 and a set of variable number tandem repeats. Clin. Microbiol. Infect. (2008) doi:10.1111/j.1469-0691.2008.02011.x.

11. Seth-Smith, H. M. B. et al. Whole-genome sequences of Chlamydia trachomatis directly from clinical samples without culture. Genome Res. (2013) doi:10.1101/gr.150037.112.

12. Christiansen, M. T. et al. Whole-genome enrichment and sequencing of Chlamydia trachomatis directly from clinical samples. BMC Infect. Dis. (2014) doi:10.1186/s12879-014-0591-3.

13. Harris, S. R. et al. Whole-genome analysis of diverse Chlamydia trachomatis strains identifies phylogenetic relationships masked by current clinical typing. Nat. Genet. (2012) doi:10.1038/ng.2214.

14. Andersson, P. et al. Chlamydia trachomatis from Australian Aboriginal people with trachoma are polyphyletic composed of multiple distinctive lineages. Nat. Commun. (2016) doi:10.1038/ncomms10688.

15. Hadfield, J. et al. Comprehensive global genome dynamics of Chlamydia trachomatis show ancient diversification followed by contemporary mixing and recent lineage expansion. Genome Res. (2017) doi:10.1101/gr.212647.116.

16. Last, A. R. et al. Population-based analysis of ocular Chlamydia trachomatis in trachoma-endemic West African communities identifies genomic markers of disease severity. Genome Med. (2018) doi:10.1186/s13073-018-0521-x.

17. Alkhidir, A. A. I. et al. Whole-genome sequencing of ocular Chlamydia trachomatis isolates from Gadarif State, Sudan. Parasites and Vectors (2019) doi:10.1186/s13071-019-3770-7.

18. Hong, K. C. et al. Lack of macrolide resistance in Chlamydia trachomatis after mass azithromycin distributions for trachoma. Emerg. Infect. Dis. (2009) doi:10.3201/eid1507.081563.

19. O’Brien, K. S. et al. Antimicrobial resistance following mass azithromycin distribution for trachoma: a systematic review. The Lancet Infectious Diseases (2019) doi:10.1016/S1473-3099(18)30444-4.

20. Keenan, J. D. et al. Azithromycin to reduce childhood mortality in sub-Saharan Africa. N. Engl. J. Med. (2018) doi:10.1056/NEJMoa1715474.

21. Bogoch, I. I., Utzinger, J., Lo, N. C. & Andrews, J. R. Antibacterial mass drug administration for child mortality reduction: Opportunities, concerns, and possible next steps. PLoS Negl. Trop. Dis. (2019) doi:10.1371/journal.pntd.0007315.

22. Bojang, A. et al. Genomic investigation of Staphylococcus aureus recovered from Gambian women and newborns following an oral dose of intra-partum azithromycin. J. Antimicrob. Chemother. (2019) doi:10.1093/jac/dkz341.

23. Marks, M. et al. Randomized Trial of Community Treatment with Azithromycin and Ivermectin Mass Drug Administration for Control of Scabies and Impetigo. Clin. Infect. Dis. (2019) doi:10.1093/cid/ciy574.

24. King, J. D. et al. Prevalence of Trachoma at Sub-District Level in Ethiopia: Determining When to Stop Mass Azithromycin Distribution. PLoS Negl. Trop. Dis. (2014) doi:10.1371/journal.pntd.0002732.

25. Thylefors, B., Dawson, C. R., Jones, B. R., West, S. K. & Taylor, H. R. A simple system for the assessment of trachoma and its complications. Bull. World Health Organ. (1987).

26. Nash, S. D. et al. Trachoma prevalence remains below threshold in five districts after stopping mass drug administration: Results of five surveillance surveys within a hyperendemic setting in Amhara, Ethiopia. Transactions of the Royal Society of Tropical Medicine and Hygiene (2018) doi:10.1093/trstmh/try096.

27. Nash, S. D. et al. Ocular Chlamydia trachomatis infection and infectious load among pre-school aged children within trachoma hyperendemic districts receiving the SAFE strategy, Amhara region, Ethiopia. PLoS Negl. Trop. Dis. 14, e0008226 (2020).

28. Moncada, J., Shayevich, C., Philip, S. S., Lucic, D. & Schachter, J. Detection of Chlamydia trachomatis and Neisseria gonorrhoeae in Rectal and Oropharyngeal Swabs and Urine Specimens from Men Who Have Sex With Men with Abbott’s M2000 RealTime. Sex. Transm. Dis. 42, 650–651 (2015).

29. Butcher, R. M. R. et al. Low Prevalence of Conjunctival Infection with Chlamydia trachomatis in a Treatment-Naïve Trachoma-Endemic Region of the Solomon Islands. PLoS Negl. Trop. Dis. (2016) doi:10.1371/journal.pntd.0004863.

30. Butcher, R. et al. Reduced-cost Chlamydia trachomatis-specific multiplex real-time PCR diagnostic assay evaluated for ocular swabs and use by trachoma research programmes. J. Microbiol. Methods (2017) doi:10.1016/j.mimet.2017.04.010.

31. Bolger, A. M., Lohse, M. & Usadel, B. Trimmomatic: A flexible trimmer for Illumina sequence data. Bioinformatics (2014) doi:10.1093/bioinformatics/btu170.

32. Langmead and Steven L Salzberg. Bowtie2. Nat. Methods (2013) doi:10.1038/nmeth.1923.Fast.

33. Li, H. et al. The Sequence Alignment/Map format and SAMtools. Bioinformatics (2009) doi: 10.1093/bioinformatics/btp352.

34. Darling, A. C. E., Mau, B., Blattner, F R. & Perna, N. T. Mauve: Multiple alignment of conserved genomic sequence with rearrangements. Genome Res. (2004) doi:10.1101/gr.2289704.

35. Edgar, R. C. MUSCLE: Multiple sequence alignment with high accuracy and high throughput. Nucleic Acids Res. (2004) doi:10.1093/nar/gkh340.

36. Stamatakis, A. RAxML version 8: A tool for phylogenetic analysis and post-analysis of large phylogenies. Bioinformatics (2014) doi:10.1093/bioinformatics/btu033.

37. Croucher, N. J. et al. Rapid phylogenetic analysis of large samples of recombinant bacterial whole genome sequences using Gubbins. Nucleic Acids Res. (2015) doi:10.1093/nar/gku1196.

38. Gouy, M., Guindon, S. & Gascuel, O. Sea view version 4: A multiplatform graphical user interface for sequence alignment and phylogenetic tree building. Mol. Biol. Evol. (2010) doi:10.1093/molbev/msp259.

39. Lutter, E. I. et al. Phylogenetic analysis of Chlamydia trachomatis tarp and correlation with clinical phenotype. Infect. Immun. (2010) doi:10.1128/IAI.00515-10.

40. Caldwell, H. D. et al. Polymorphisms in Chlamydia trachomatis tryptophan synthase genes differentiate between genital and ocular isolates. J. Clin. Invest. (2003) doi:10.1172/JCI17993.

41. Gomes, J. P. et al. Polymorphisms in the nine polymorphic membrane proteins of Chlamydia trachomatis across all serovars: Evidence for serovar da recombination and correlation with tissue tropism. J. Bacteriol. (2006) doi: 10.1128/JB.188.1.275-286.2006.

42. West, S. K. Milestones in the fight to eliminate trachoma. Ophthalmic and Physiological Optics (2020) doi:10.1111/opo.12666.

43. S.E., S. et al. Single dose mass drug administration of azithromycin decreases malaria incidence in a large cohort treated for ocular trachoma. Am. J. Trop. Med. Hyg. (2011).

44. Schachterle, S. E. et al. Short-term malaria reduction by single-dose azithromycin during mass drug administration for trachoma, Tanzania. Emerg. Infect. Dis. (2014) doi:10.3201/eid2006.131302.

45. Harrison, M. A. et al. Impact of mass drug administration of azithromycin for trachoma elimination on prevalence and azithromycin resistance of genital Mycoplasma genitalium infection. Sex. Transm. Infect. 95, 522–528 (2019).

46. Coles, C. L. et al. Association of mass treatment with azithromycin in trachoma-endemic communities with short-term reduced risk of diarrhea in young children. Am. J. Trop. Med. Hyg. (2011) doi:10.4269/ajtmh.2011.11-0046.

47. Whitty, C. J. M., Glasgow, K. W., Sadiq, S. T., Mabey, D. C. & Bailey, R. Impact of community-based mass treatment for trachoma with oral azithromycin on general morbidity in Gambian children. Pediatr. Infect. Dis. J. (1999) doi:10.1097/00006454-199911000-00003.

48. Keenan, J. D. et al. Childhood mortality in a cohort treated with mass azithromycin for trachoma. Clin. Infect. Dis. (2011) doi:10.1093/cid/cir069.

49. See, C. W. et al. The effect of mass azithromycin distribution on childhood mortality: Beliefs and estimates of efficacy. Am. J. Trop. Med. Hyg. (2015) doi:10.4269/ajtmh.15-0106.

50. Keenan, J. D. et al. Longer-term assessment of azithromycin for reducing childhood mortality in Africa. N. Engl. J. Med. (2019) doi:10.1056/NEJMoa1817213.

51. Mack, I. et al. Antimicrobial Resistance Following Azithromycin Mass Drug Administration: Potential Surveillance Strategies to Assess Public Health Impact. Clin. Infect. Dis. (2020) doi:10.1093/cid/ciz893.

52. Keenan, J. D. et al. Mass azithromycin distribution for hyperendemic trachoma following a cluster-randomized trial: A continuation study of randomly reassigned subclusters (TANA II). PLoS Med. (2018) doi:10.1371/journal.pmed.1002633.

53. Bojang, E. et al. Short-term increase in prevalence of nasopharyngeal carriage of macrolide-resistant Staphylococcus aureus following mass drug administration with azithromycin for trachoma control. BMC Microbiol. (2017) doi:10.1186/s12866-017-0982-x.

54. Coles, C. L. et al. Mass distribution of azithromycin for trachoma control is associated with increased risk of azithromycin-resistant streptococcus pneumoniae carriage in young children 6 months after treatment. Clin. Infect. Dis. (2013) doi:10.1093/cid/cit137.

55. Skalet, A. H. et al. Antibiotic selection pressure and macrolide resistance in Nasopharyngeal Streptococcus pneumoniae: A cluster-randomized clinical trial. PLoS Med. (2010) doi:10.1371/journal.pmed.1000377.

56. Abeles, S. R. et al. Microbial diversity in individuals and their household contacts following typical antibiotic courses. Microbiome (2016) doi:10.1186/s40168-016-0187-9.

57. Parker, E. P. K. et al. Changes in the intestinal microbiota following the administration of azithromycin in a randomised placebo-controlled trial among infants in south India. Sci. Rep. (2017) doi:10.1038/s41598-017-06862-0.

58. Doan, T. et al. Mass azithromycin distribution and community microbiome: A cluster-randomized trial. Open Forum Infect. Dis. (2018) doi:10.1093/ofid/ofy182.

59. Doan, T. et al. Gut microbiome alteration in MORDOR I: a community-randomized trial of mass azithromycin distribution. Nature Medicine (2019) doi:10.1038/s41591-019-0533-0.

60. J.C., S. et al. Increased resistance to azithromycin in e. coli following mass treatment for trachoma control in rural tanzania. Am. J. Trop. Med. Hyg. (2012).

61. Seidman, J. C. et al. Increased carriage of macrolide-resistant fecal E. coli following mass distribution of azithromycin for trachoma control. Int. J. Epidemiol. (2014) doi:10.1093/ije/dyu062.

62. Dean, D., Rothschild, J., Ruettger, A., Kandel, R. P. & Sachse, K. Zoonotic Chlamydiaceae species associated with trachoma, Nepal. Emerg. Infect. Dis. (2013) doi:10.3201/eid1912.130656.

63. Ghasemian, E. et al. Detection of chlamydiaceae and chlamydia-like organisms on the ocular surface of children and adults from a trachoma-endemic region. Sci. Rep. (2018) doi:10.1038/s41598-018-23887-1.

64. Burton, M. J. et al. What Is causing active trachoma? The role of nonchlamydial bacterial pathogens in a low prevalence setting. Investig. Ophthalmol. Vis. Sci. (2011) doi:10.1167/iovs.11-7326.

65. Hu, V H. et al. Bacterial infection in scarring trachoma. Investig. Ophthalmol. Vis. Sci. (2011) doi:10.1167/iovs.10-5829.

66. Burr, S. E. et al. Association between Ocular Bacterial Carriage and Follicular Trachoma Following Mass Azithromycin Distribution in The Gambia. PLoS Negl. Trop. Dis. (2013) doi:10.1371/journal.pntd.0002347.

67. Zhou, Y. et al. The conjunctival microbiome in health and trachomatous disease: A case control study. Genome Med. (2014) doi:10.1186/s13073-014-0099-x.

68. Hu, V. H. et al. Non-chlamydial bacterial infection and progression of conjunctival scarring in trachoma. Investig. Ophthalmol. Vis. Sci. (2018) doi:10.1167/iovs.17-23381.

69. Pickering, H. et al. Conjunctival Microbiome-Host Responses Are Associated With Impaired Epithelial Cell Health in Both Early and Late Stages of Trachoma. Front. Cell. Infect. Microbiol. (2019) doi:10.3389/fcimb.2019.00297.

70. Kari, L. et al. Pathogenic Diversity among Chlamydia trachomatis Ocular Strains in Nonhuman Primates Is Affected by Subtle Genomic Variations. J. Infect. Dis. (2008) doi:10.1086/525285.

71. Brunelle, B. W. & Sensabaugh, G. F. The ompA gene in Chlamydia trachomatis differs in phylogeny and rate of evolution from other regions of the genome. Infect. Immun. (2006) doi: 10.1128/IAI.74.1.578-585.2006.

72. Pickering, H. et al. Genome-wide profiling of humoral immunity and pathogen genes under selection identifies immune evasion tactics of Chlamydia trachomatis during ocular infection. Sci. Rep. (2017) doi:10.1038/s41598-017-09193-2.

73. Sun, G. et al. Structural and functional analyses of the major outer membrane protein of Chlamydia trachomatis. J. Bacteriol. (2007) doi:10.1128/JB.00552-07.

74. Sauvage, E., Kerff, F., Terrak, M., Ayala, J. A. & Charlier, P. The penicillin-binding proteins: Structure and role in peptidoglycan biosynthesis. FEMS Microbiology Reviews (2008) doi:10.1111/j.1574-6976.2008.00105.x.

75. Saini, A., Mapolelo, D. T., Chahal, H. K., Johnson, M. K. & Outten, F. W. SufD and SufC ATPase activity are required for iron acquisition during in vivo Fe-S cluster formation on SufB. Biochemistry (2010) doi:10.1021/bi1011546.

76. Zhang, J., Lietman, T., Olinger, L., Miao, Y. & Stephens, R. S. Genetic diversity of Chlamydia trachomatis and the precalence of trachoma. Pediatr. Infect. Dis. J. (2004) doi:10.1097/01.inf.0000115501.60397.a6.

77. Chin, S. A. et al. Diversity of chlamydia trachomatis in trachoma-hyperendemic communities treated with azithromycin. Am. J. Epidemiol. (2018) doi:10.1093/aje/kwy071.

78. Dawson, C., Wood, T. R., Rose, L. & Hanna, L. Experimental Inclusion Conjunctivitis in Man: III. Keratitis and Other Complications. Arch. Ophthalmol. (1967) doi:10.1001/archopht.1967.00980030343015.

79. Tarizzo, M. L., Nataf, R. & Nabli, B. Experimental inoculation of thirteen volunteers with agent isolated from inclusion conjunctivitis. Am. J. Ophthalmol. (1967) doi:10.1016/0002-9394(67)94093-7.

80. Taylor, H. R. Development of immunity to ocular chlamydial infection. Am. J. Trop. Med. Hyg. (1990) doi:10.4269/ajtmh.1990.42.358.

