## Supplementary Information for "Genomics of Ocular *Chlamydia trachomatis* after 5 years of SAFE interventions for trachoma in Amhara, Ethiopia"

SI Figure 1.

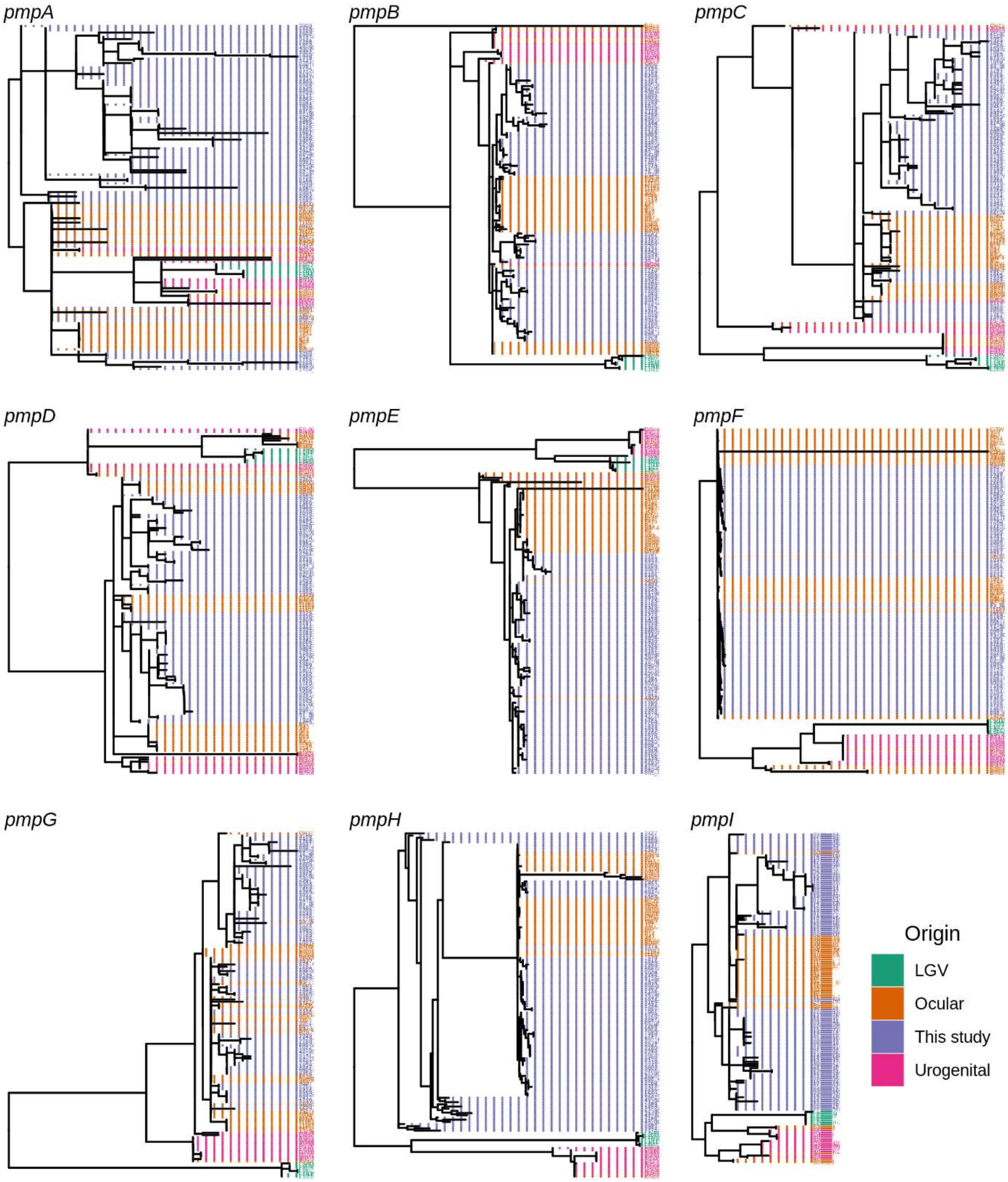


Maximum likelihood reconstruction of phylogeny of the polymorphic membrane proteins (pmps) of ocular *Chlamydia trachomatis* (*Ct*) sequences from Amhara, Ethiopia. Phylogeny of *ompA* from 99 *Ct* sequences from Amhara and 183 *Ct* clinical and reference strains. Amharan *Ct* sequences were mapped to *Ct* A/HAR-13 using Bowtie2. SNPs were called using SAMtools/BCFtools. Phylogenies were computed with RAxML from a variable sites alignment using a GTR + gamma model and are midpoint rooted. *Ct* sequences are coloured by country of origin of the sample ("Origin").

SI Figure 2.

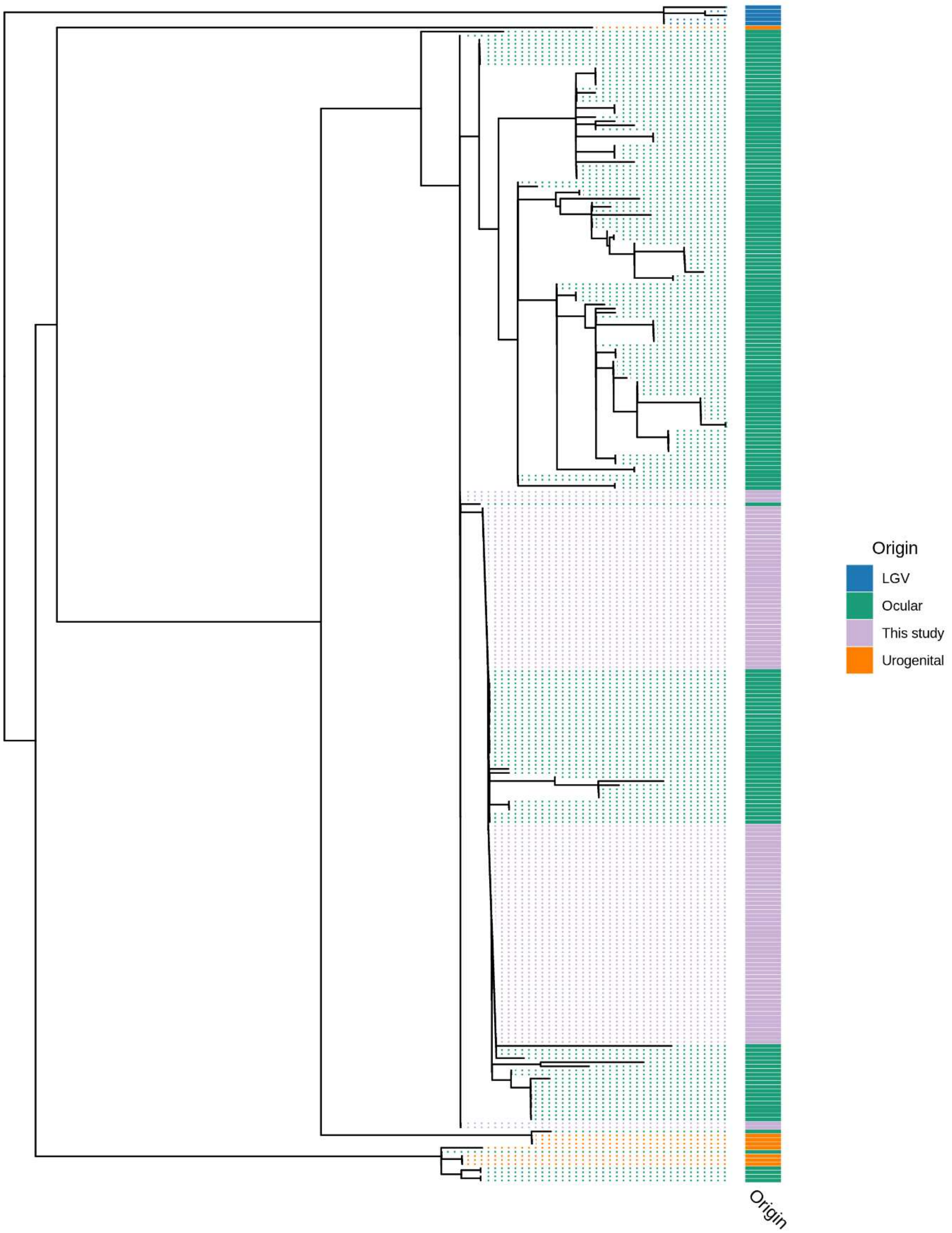


Maximum likelihood reconstruction of plasmid phylogeny of ocular *Chlamydia trachomatis* (*Ct*) sequences from Amhara, Ethiopia. Phylogeny of *ompA* from 99 *Ct* sequences from Amhara and 183 *Ct* clinical and reference strains. Ethiopian *Ct* sequences were mapped to *Ct* A/HAR-13 using Bowtie2. SNPs were called using SAMtools/BCFtools. Phylogenies were computed with RAxML from a variable sites alignment using a GTR + gamma model and are midpoint rooted. *Ct* sequences are coloured by country of origin of the sample (“Origin”).

SI Figure 3.

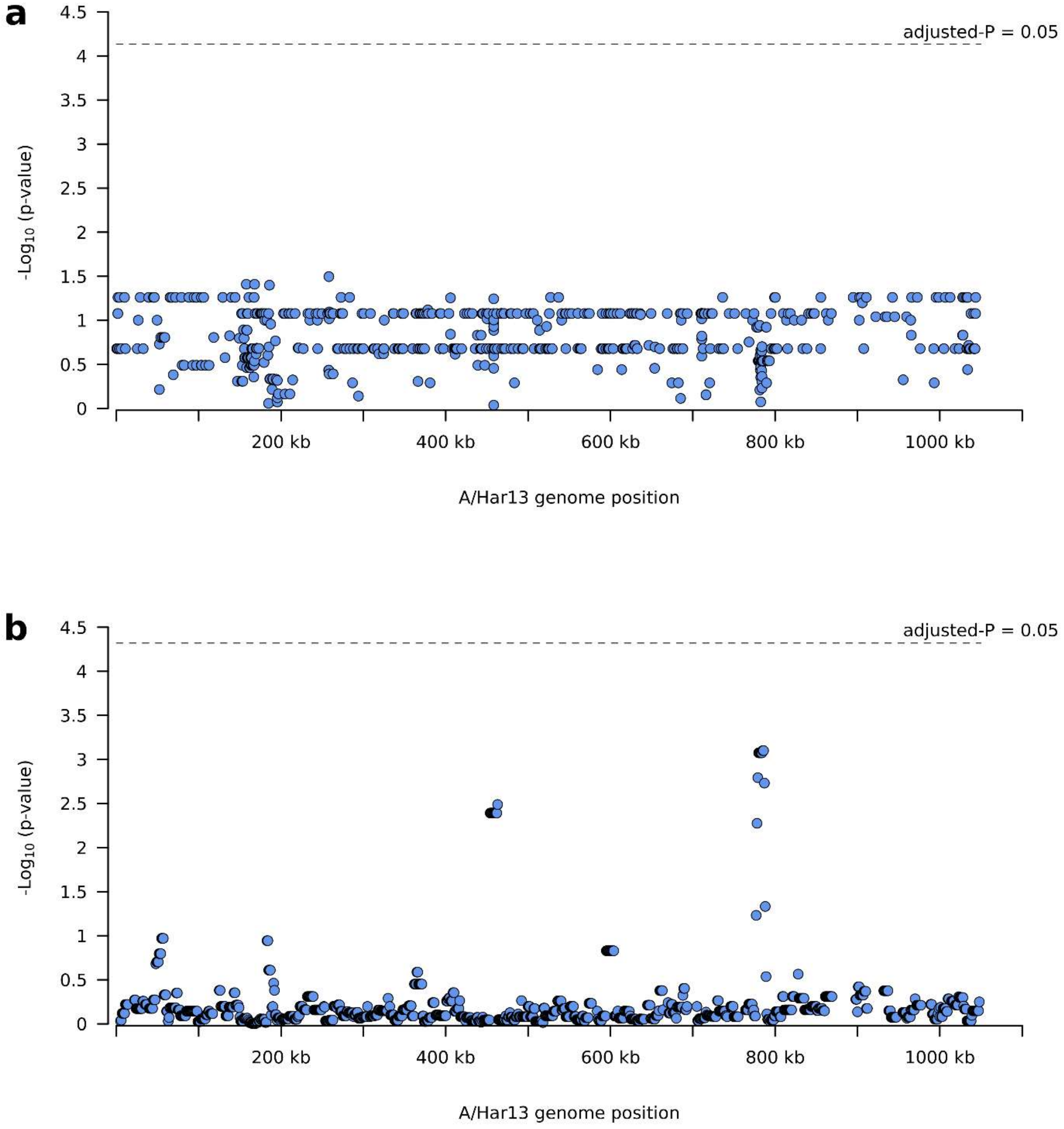


Polymorphisms on the *Chlamydia trachomatis* (*Ct*) genome associated with village-level *Ct* infection prevalence. a) No single nucleotide polymorphisms were significantly associated with village-level *Ct* infection prevalence. b) No polymorphic regions were significantly associated with village-level *Ct* infection prevalence.

SI Figure 4.

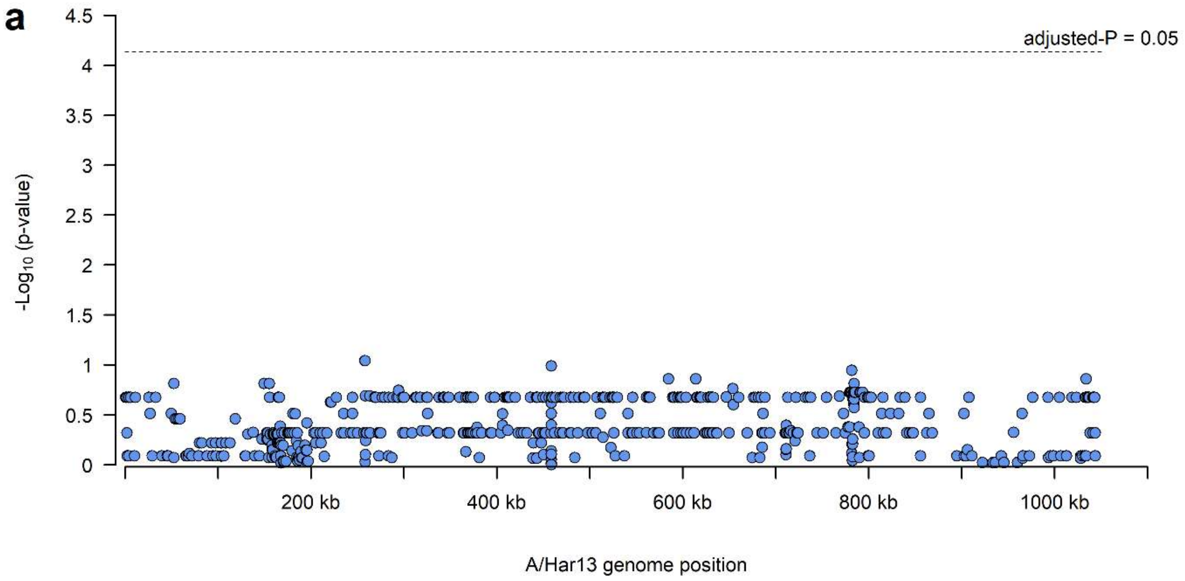


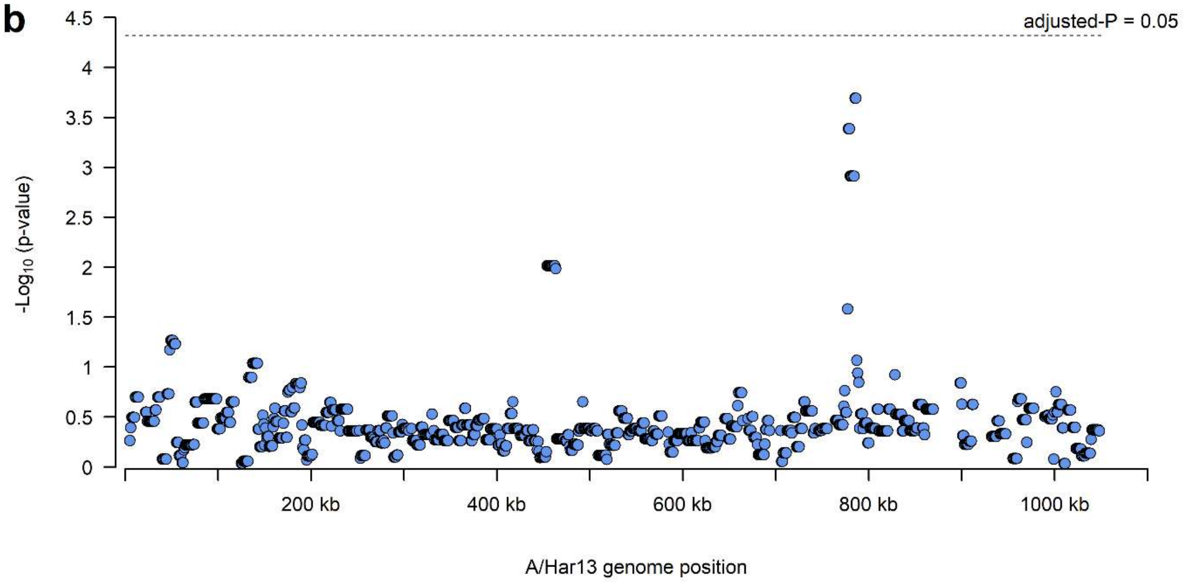


Polymorphisms on the *Chlamydia trachomatis* (*Ct*) genome associated with village-level TI prevalence. a) No single nucleotide polymorphisms were significantly associated with village-level TI prevalence. b) No polymorphic regions were significantly associated with village-level TI prevalence.

SI Figure 5.

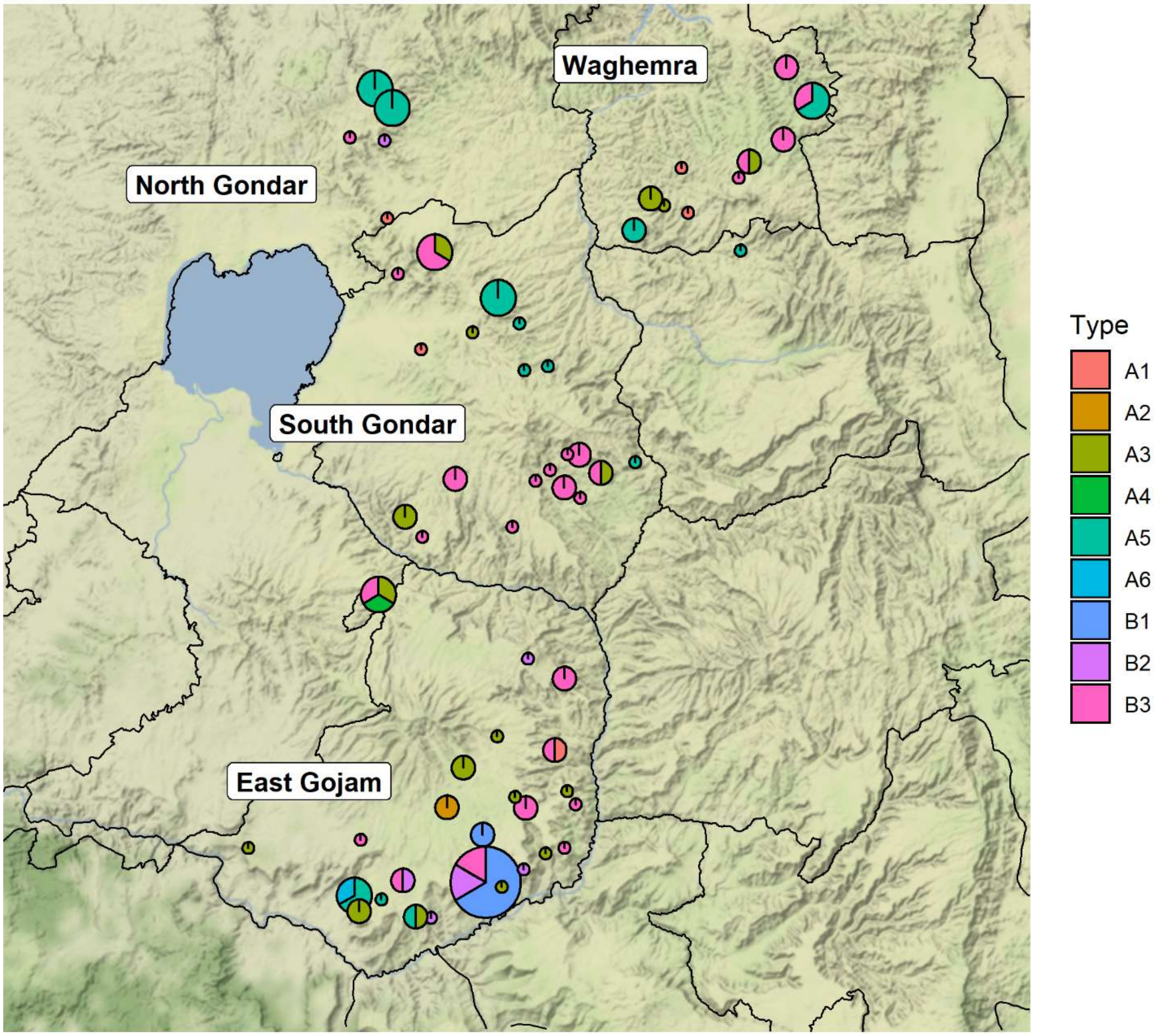


Geographical distribution of *ompA* types. Four zones in Amhara, Ethiopia were represented in this study. Pie charts represent village-level *Ct* prevalence (pie diameter) and presence of *ompA* types. Maps were generated using R package ggmap, shape files were obtained from Google Maps.
